# The Good, the Bad and the Stochastic: How Living in Groups Innately Supports Cooperation

**DOI:** 10.1101/2021.02.21.431661

**Authors:** Klaus F. Steiner

## Abstract

Based on theoretical considerations and computer simulations, I show that living in groups brings advantages for cooperative traits through purely stochastic effects that result from the division of a population into groups. These advantages can be sufficient to compensate individual selection pressures that may be associated with the cooperative traits. In more complex agent-based simulation models, this effect combined with some migration between the groups leads to stable dynamic equilibria between cooperative and defective replicators in the population.

## Introduction

A central question in evolutionary biology since Charles Darwin is how cooperative traits emerge and prevail, although they can entail costs and disadvantages for the cooperative actors. However, countless examples in biology, from bacterial colonies to insect states and human societies, demonstrate that there must be ways and means for the evolution of traits that are “for the good of the group”.

Nowak (2012) lists five mechanisms for the evolution of cooperation: direct and indirect reciprocity as well as spatial selection, multi-level selection, and kin selection. A rather controversial mechanism is multi-level selection, an enhanced group selection, which aims to explain that there are different levels on which natural selection works. Some concepts and models show that group and multi-level selection might work (Wilson 1975, Traulsen & Nowak 2006, Wilson&Wilson 2008), but there are also strong opposing views.

Maynard Smith & Szathmáry (1996) argue that in the “Trait Group Model” (Wilson 1975) the frequency of altruistic replicators can only increase if altruistic replicators come together non-randomly or if there are synergistic effects on fitness (Fig. 1). Representatives of group and multi-level selection, however, argue that competition between groups would be necessary that group selection takes place (Wilson&Wilson 2008).

**Figure 1:**
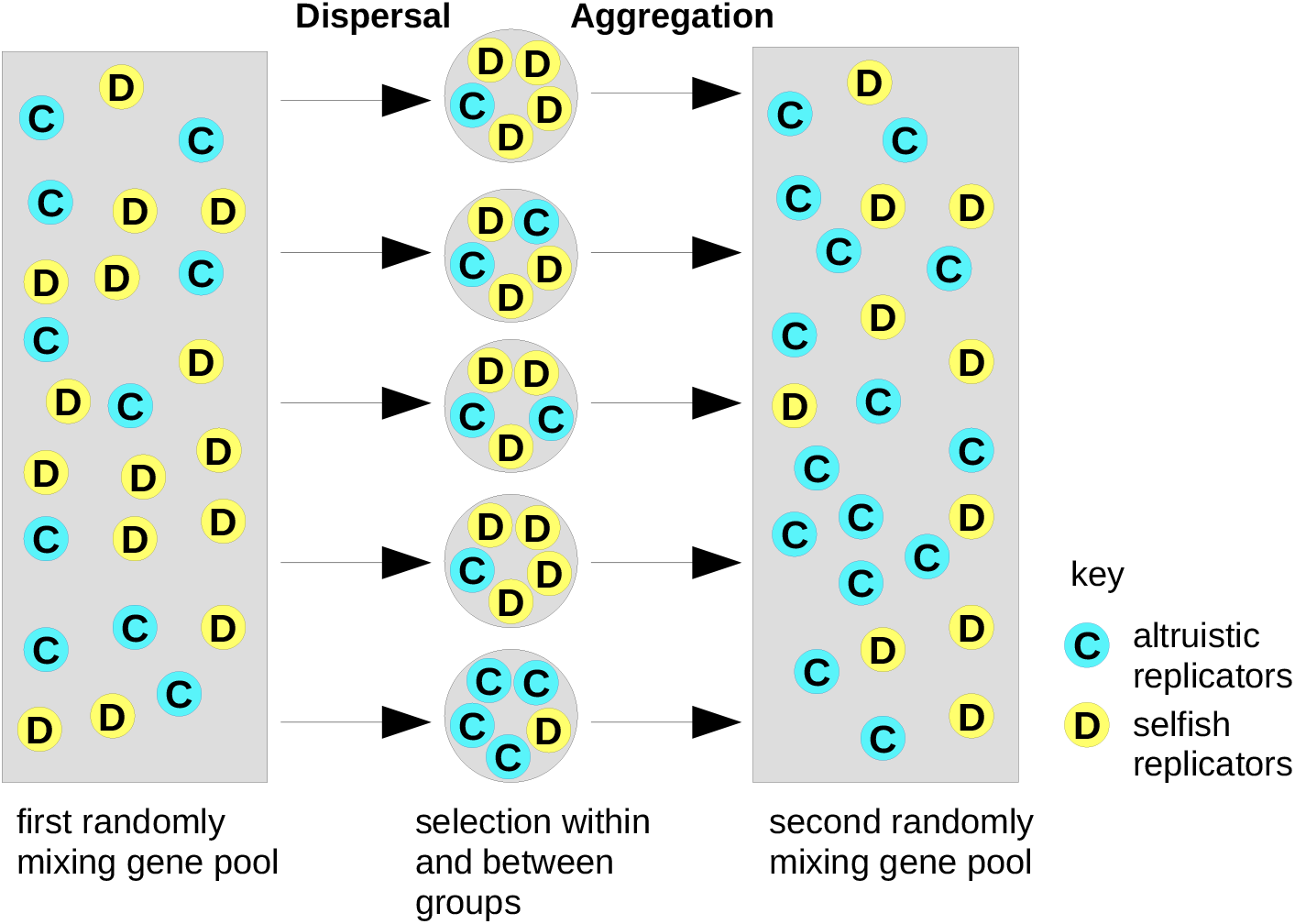
Description and criticism of Wilson’s “trait group” model from Maynard Smith and Szathmáry (1996), they wrote: “If there is to be an increase in the frequency of altruistic replicators, as shown, then either (1) altruistic replicators must assort together non-randomly or (2) there must be synergistic fitness effects, so that a group of altruistic replicators contributes disproportionately to the next generation.”

Based on some calculations and multi-agent simulations, however, I come to the conclusion that cooperative features alone achieve advantages through the division of a population into groups and through purely stochastic effects that occur. These effects can be sufficiently large to compensate intrinsic selection pressures that may be associated with the cooperative trait.

As a further consequence, under suitable circumstances, the migration of individuals leads to stable dynamic equilibria between cooperative (***C***) and defective (***D***) traits in the population.

## About the stochastic advantage of being cooperative in the group

The basic assumption of the following considerations is a population of cooperative individuals with trait ***C*** and selfish defectors with trait ***D***, who are divided into random groups. The cooperators each provide the contribution *b=1* to their group, the contribution of defectors is *b=0*. The sum of contributions is divided equally in the group, regardless of whether it is ***D*** or ***C***. Consequently in a group with only ***C***, each member receives the individual benefit *B*_*i*_*=1*, in a ***D*** only group there is nothing to distribute, the benefit is *B*_*i*_*=0*, and in mixed groups the individual benefit payoff lies in between.

For example, Fig. 2 shows a population of 5 cooperators and 5 defectors (written as 5C5D for short). As a single group there are thus 5 contributions from ***C*** and none from ***D***, the overall benefit is therefore 5. Divided by 10, each individual receives the benefit *B*_*i*_*=0*.*5*. The total benefit for each trait is therefore *B*_*C*_*=B*_*D*_*=2*.*5* for both the 5 ***C*** and the 5 ***D***.

**Figure 2:**
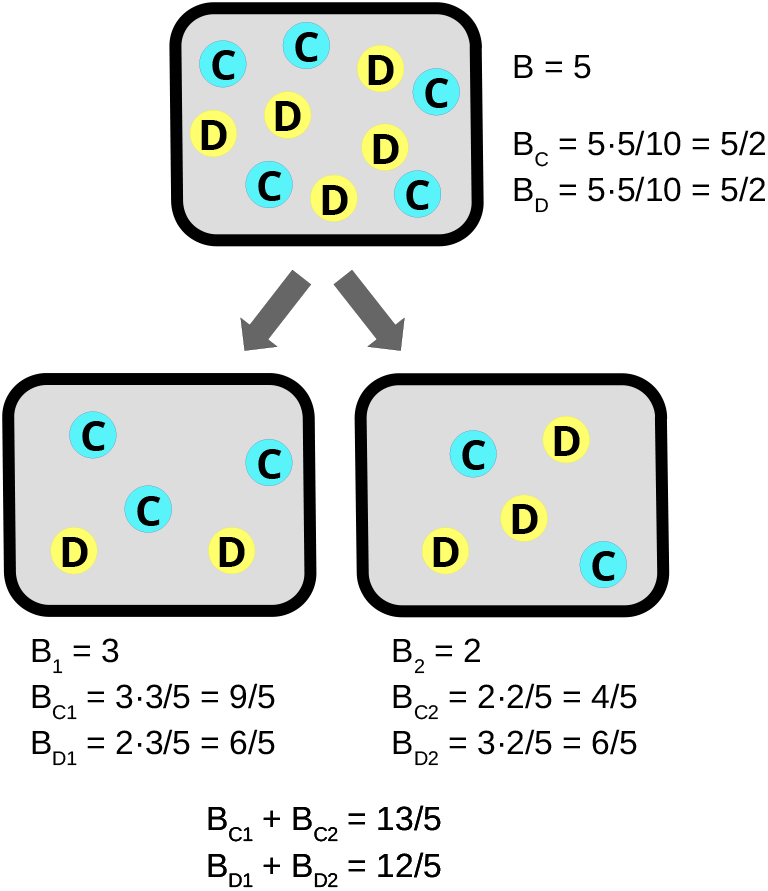
Distribution of the group benefits for **C** and **D**. After splitting into two groups, the total benefit for trait **C** is greater and for **D** less than before.

Let us now divide the 5C5D population into two groups, for example into the most likely combination 3C2D and 2C3D. The group 3C2D achieves the group benefit *B*_*1*_*=3* through the 3 cooperators. Divided among the five members, each individual in this group receives *B*_*i*_*=3/5*. The other group (2C3D) has two cooperators therefore the group benefit is only *B*_*2*_*=2*, which is *B*_*i*_*=2/5* per member.

The total benefit for trait ***C*** is *B*_*C1*_*=3·3/5* from group 3C2D and *B*_*C2*_*=*2·2/5 from group 2C3D, i.e. 3·3/5 + 2·2/5 = 13/5.

In contrast, the overall benefit for ***D*** is *B*_*D1*_*=2·3/5* from group 3C2D and *B*_*D2*_*=3·2/5* from group 2C3D, so 2·3/5 + 3·2/5 = 12/5.

This means that simply by dividing the population in two groups, the overall benefit for trait ***C*** has increased from 5/2 to 13/5 and that for ***D*** has decreased from 5/2 to 12/5. Specifically, the factors are 1.04 or +4% for ***C*** and 0.96 or −4% for ***D*** related to the population 5C5D. The absolute increase of the mean benefit for each of the five ***C***-individuals is 0.02, from 0.5 to 0.52. Conversely, the loss for every ***D*** is also 0.02, from 0.5 to 0.48.

Figure 3 shows the benefit factors for a number of grouping combinations for two differently composed populations of 10 individuals, respectively. It is obvious that the more the ***C:D*** ratio in the group deviates from that in the population, the more the factors deviate from one. The benefit payoff can even be a multiple for ***C*** and it can also be completely eliminated for ***D***, as in the case of 3C0D: 0C7D, where the trait ***C*** has a benefit of three. Compared to 9/10 in the undivided 3C7D population, this gives a factor of 3.333. However, ***D*** does not achieve any benefit, since it only occurs in the ***D-***only group.

**Figure 3:**
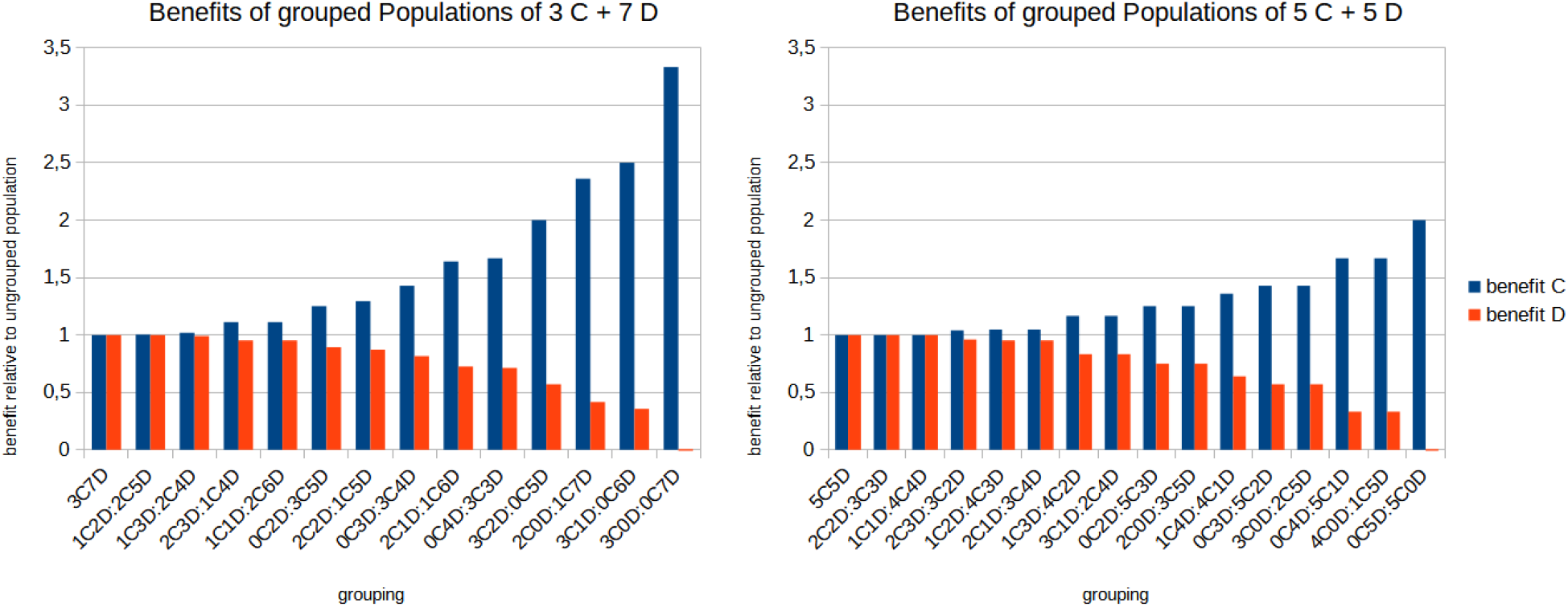
Calculated benefit factors for traits **C** and **D** and different group divisions compared to the undivided population. Details in text.

In addition, one can see that through any grouping, trait ***C*** never has a disadvantage and ***D*** never has an advantage. Both can achieve a factor of 1 only if the groups have the same symmetry (***C:D***-ratios) as the population (e.g. the two groups 2C2D: 3C3D related to 5C5D). Further simulated results for 16 groups with different group sizes, ***C:D*** ratios and group variability are shown in Fig. 4. These simulation results also show that the benefit shifts primarily depend on the variability of the groups, and also that the difference (*Δ)* between ***C*** and ***D*** is independent of the ratio between ***C*** and ***D***. This *Δ* or total benefit shift also defines the limits for the possible additional costs for ***C***, which could be compensated by this grouping effect.

**Figure 4:**
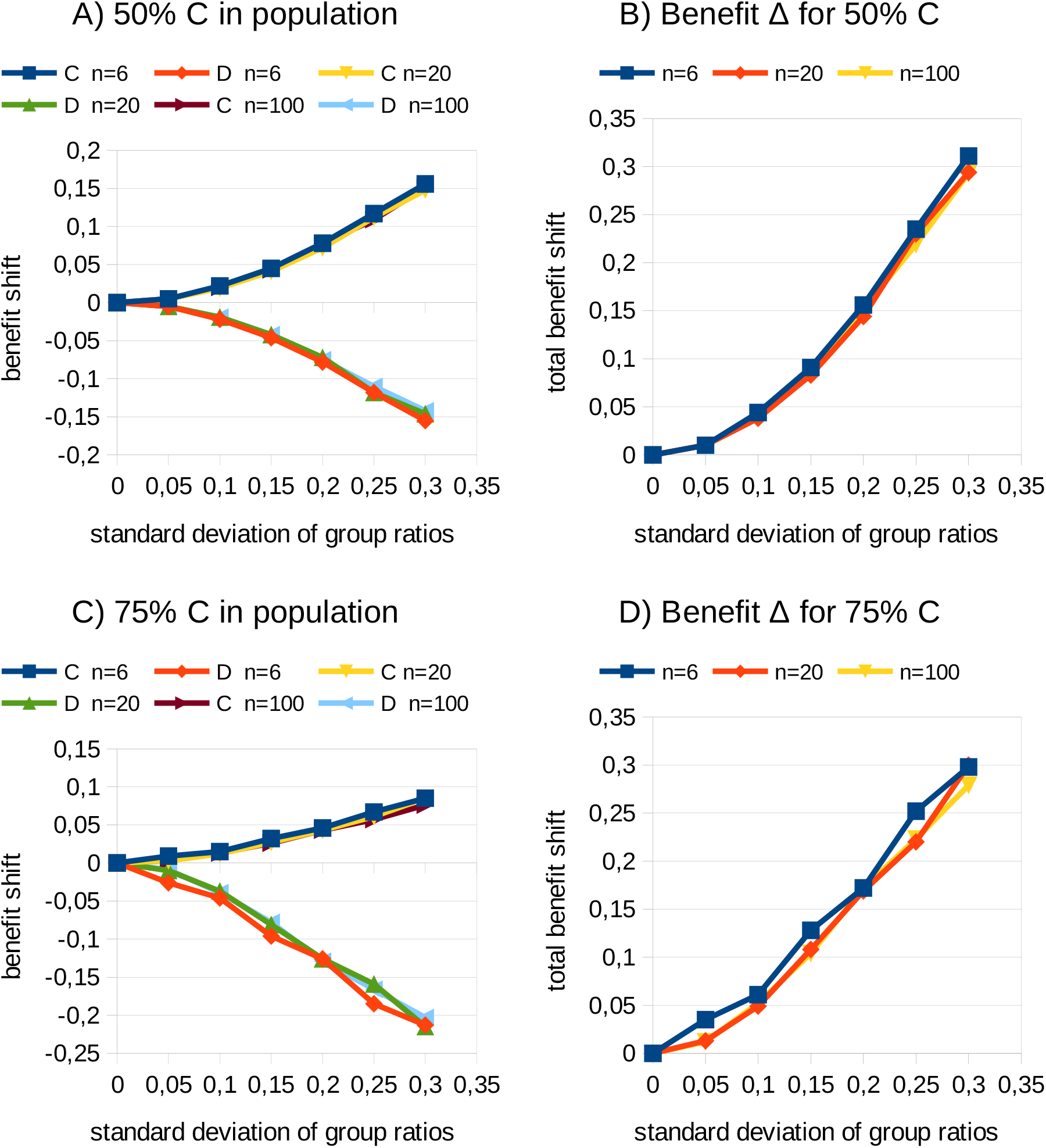
Benefit shifts and the variability of groups. Simulated populations consisted of 16 groups with n=6, 20 and 100 members respectively. The abscissas show the dispersion of the groups as the standard deviation of the group ratios and were examined from 0 to 0.3 in steps of 0.05. A and B show the results for a population with an average of 50% cooperators and 50% defectors (i.e. mean of group ratios = 0.5), in plots C and D the populations consisted on average of 75 % cooperators and 25 % defectors (mean of group ratios = 0.75). In A and C, the results for cooperators and defectors are plotted separately, B and D show the difference (Δ) between them, i.e. the entire benefit shift between the traits.

Since this effect occurs with every split into smaller groups, ***C*** should tend towards smaller groups, and ***D*** towards larger groups. This is likely to be counteracted by pressures from the advantages and disadvantages of group size (“group effects”). However, I will not go into further detail here, but rather explicitly define the maximum group sizes in my models. Up to this specified size, I treat all groups equally, provided they consist of at least two members.

## From grouping effect to system dynamics

I will demonstrate how these advantages from the group formation for cooperators (***C***) and the disadvantages for defectors (***D***) affect the dynamics of the trait distribution in the population by using increasingly complex simulation models.

### When the payoff equals the odds to survive and survivors replicate

In my simulation models, the benefit is transferred directly to the probability of survival or mortality. For a group of defectors (***D***) only, this means that every group member is eliminated in the next step. In a ***C***-only group, everyone survives. In a mixed group like C3D2, each member survives with *p=3/5* (mortality is *p=2/5*), in a group C2D7 it would be *p=2/9* (and mortality *p=7/9*). Under this assumption, the number of ***C*** in the (full) group is always equal to the number of survivors. But whoever survives is random and independent of trait ***C*** or ***D***.

In the following multi-agent simulations, there are 16 groups consisting of *n* randomly set replicators ***C*** and ***D***. In each step (= generation) the replicators are eliminated with the probability described above. Then the remaining group members replicate proportionally and inherit their trait ***C*** or ***D*** until their group is full again, i.e. there are 5 members. Finally group mortality comes back into play and so on.

Fig. 5 shows a typical development of such a population in a multi-agent simulation. After less than 10 generations, all defectors disappear and there are only groups with cooperators as well as extinct groups.

**Figure 5:**
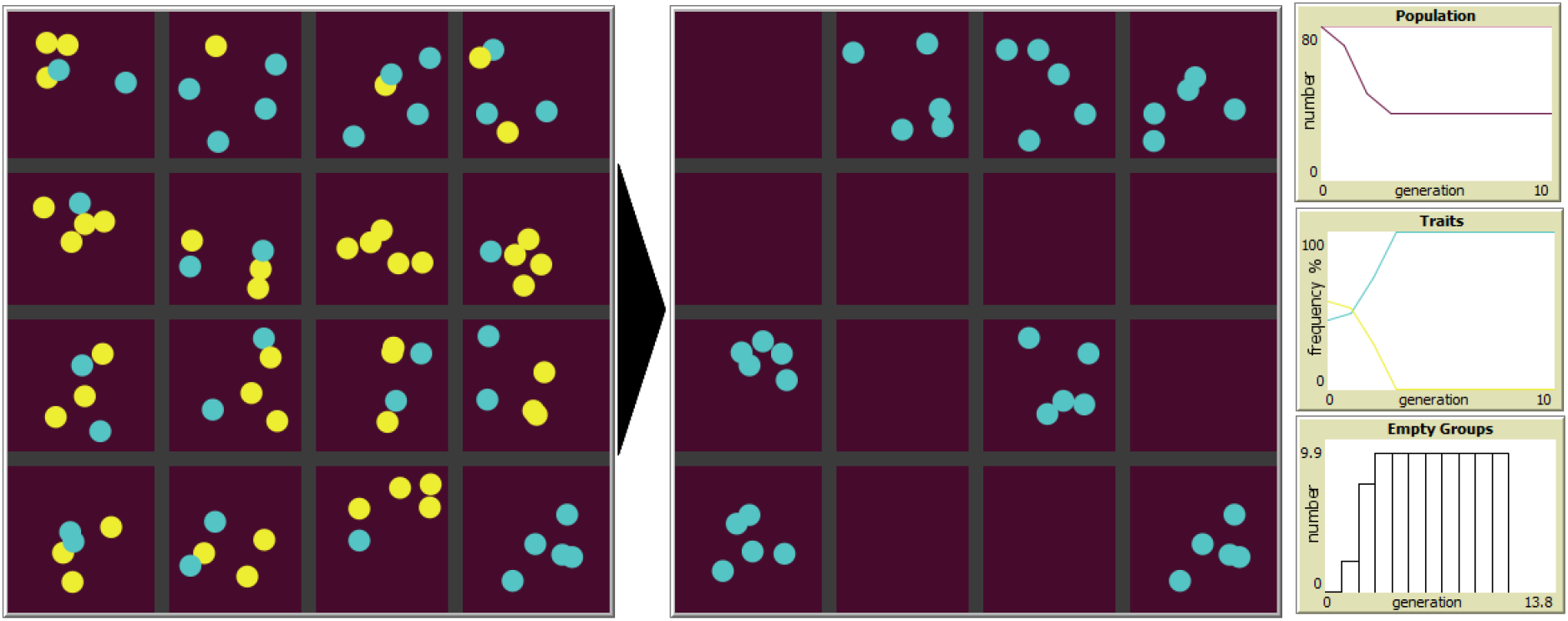
Development over 10 generations without migration: there are only cooperators left, all defectors are extinct and about half of the groups are extinguished. Left: At the beginning with random distribution. Right: After 10 generations there are only cooperators and extinct groups. Cooperators **C** blue, defectors **D** yellow; 16 groups with 5 individuals each. The diagram right above: Population size in each generation; middle: traits plot shows the relative frequency of **C** (blue line) and **D** (yellow) over the generations; bottom plot shows the number of empty groups in each generation. This and the following simulation models I have created in the modeling environment Netlogo (Wilensky, 1999).

The groups are firmly defined in my models and best understood as territories that are appropriately occupied or not. If empty or not fully occupied groups are allowed to be populated with migrating offspring from full groups, then all groups are occupied with cooperators ***C*** within a few generations (see Fig. 6).

**Figure 6:**
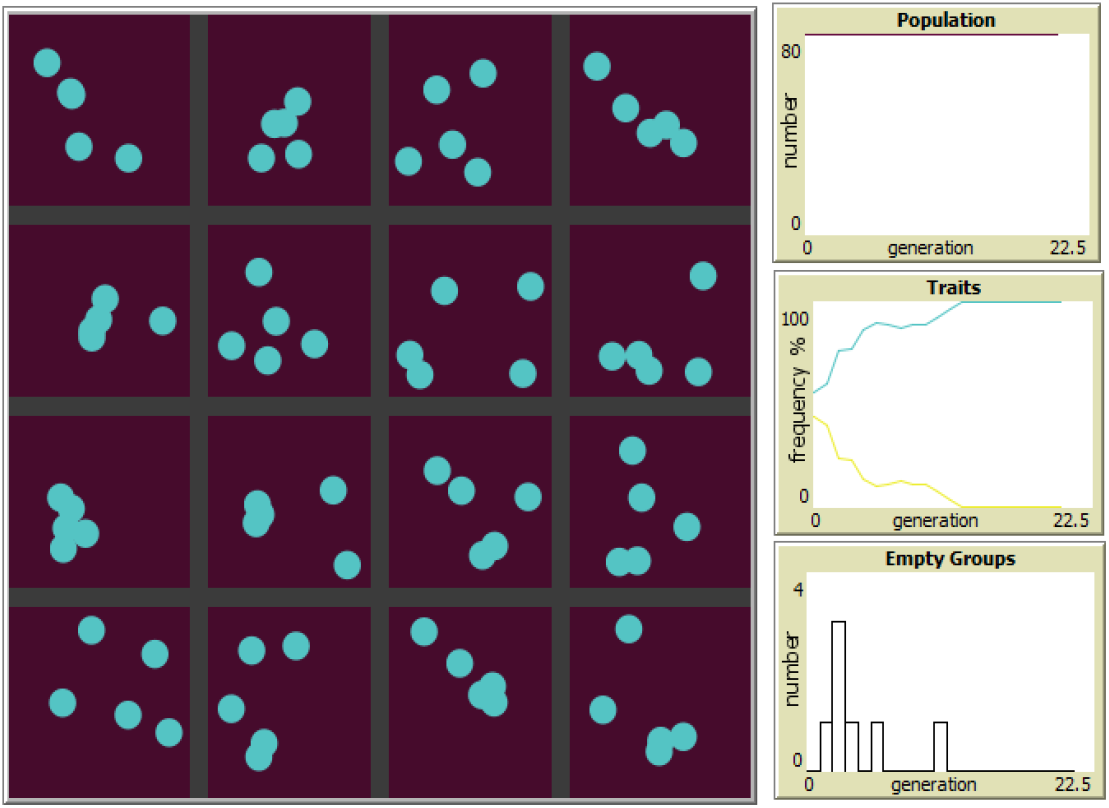
Development of a random distribution after 20 generations with migration results in a population of cooperators **C** only, all groups are full and all **D** are extincted. Cooperators C blue, defectors D yellow; 16 groups with a maximum of 5 individuals each. The diagram above: Population is maximum over the time; middle: Traits plot shows increasing frequency of **C** (blue line) and decreasing **D** (yellow) until only **C** remains; bottom plot shows the number of empty groups in each generation

### Models with mutation and extra costs for cooperators

I also considered mutation for more realistic situations. Thus, with a certain probability *m*, a descendant has a different trait than its producer, i.e. ***C*** instead of ***D*** or vice versa. As a result, a trait alone cannot be eradicated. In the simulations, a dynamic equilibrium between ***C*** and ***D*** is usually established.

In nature, cooperative traits result in additional costs for the individual, since cooperation usually entails a certain additional effort or risk for the individual. Thus, for more realistic simulations, I introduced different selection pressures as intrinsic mortality rates on the two traits, whereby a higher mortality rate on ***C*** is of course the more interesting and realistic case. I incorporated this into the simulations in the form of different intrinsic mortality rates for ***C*** and ***D***. These intrinsic mortality rates are completely independent of the group or other influences.

The process is now as follows: The individuals surviving the “group selection” (according to the mortality rate due to their group’s benefit payoff, which leads to the stochastic effect described above) and “individual selection” (corresponding to the intrinsic mortality rates of their traits) each produce asexually *N* offspring with a mutation rate of *m*. The offspring first fill up their own group, and if it is full, they migrate to empty or not yet full groups. When all groups are full, the excess offspring are eliminated. Then the cycle starts all over again with the selection. In some simulations I also add random migration between the groups at given rates.

As a typical result for a wide range of parameters, stable equilibria are established with this model. In addition to the stochastic grouping effect, the flow of offspring from the more prosperous groups with mainly ***C***, supports the cooperators. For example, about 90% of the population in Fig. 7 consist of cooperators, although their intrinsic mortality is 5% higher than that of the defectors.

**Figure 7:**
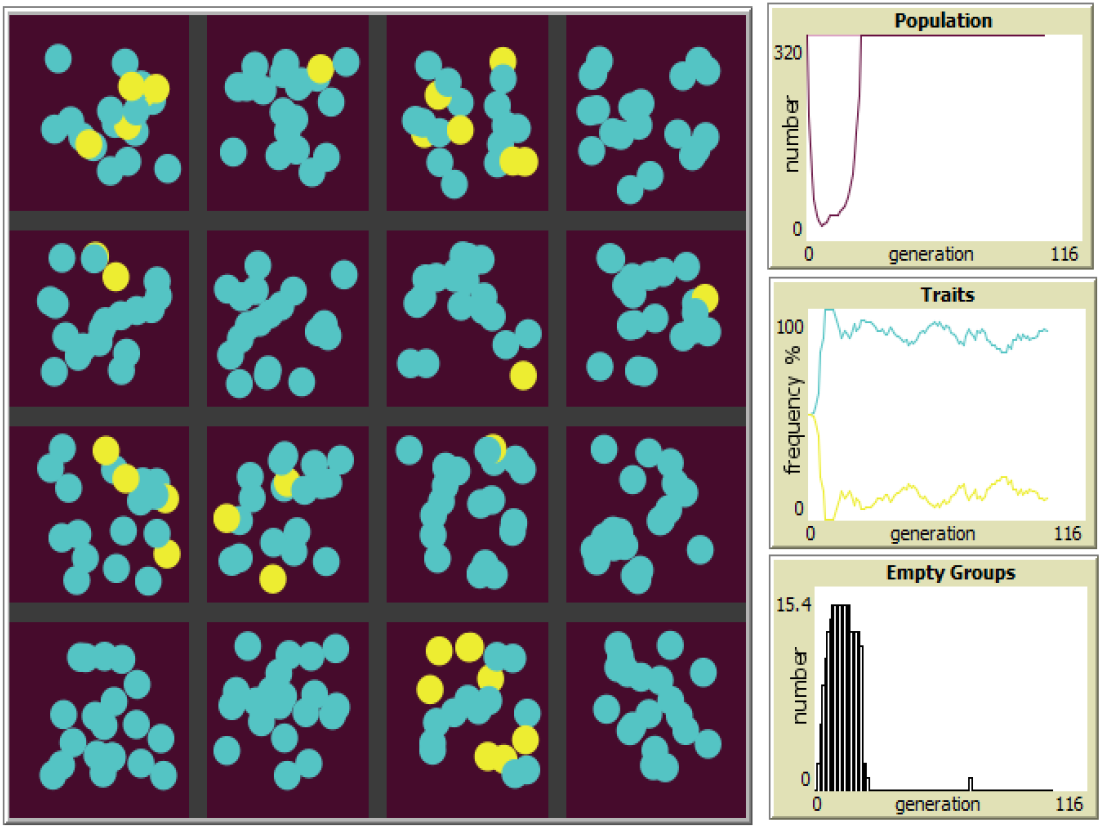
Stable steady state with mutation and different selection pressures over 100 generations. Cooperators C blue, defectors D yellow. 16 groups with a maximum of 20 individuals each. Offspring per survivor N=1; Mutation rate m=0.02; Mortality for C 25%, for D 20%. The traits diagram shows the relatively stable distribution of almost 90% C (blue) and about 10% D (yellow) over the generations. The population decreases sharply after the start with the random population, but soon reaches the maximum size again and then remains at this level.

If values are too extreme, such as very different or high mortality rates for ***C*** and ***D***, the whole population quickly collapses and dies out. Values in the marginal area, however, can lead to regular to chaotic fluctuations in the distribution of ***C*** and ***D*** as well as in population size, which can persist over hundreds or even thousands of generations (Fig. 8).

**Figure 8:**
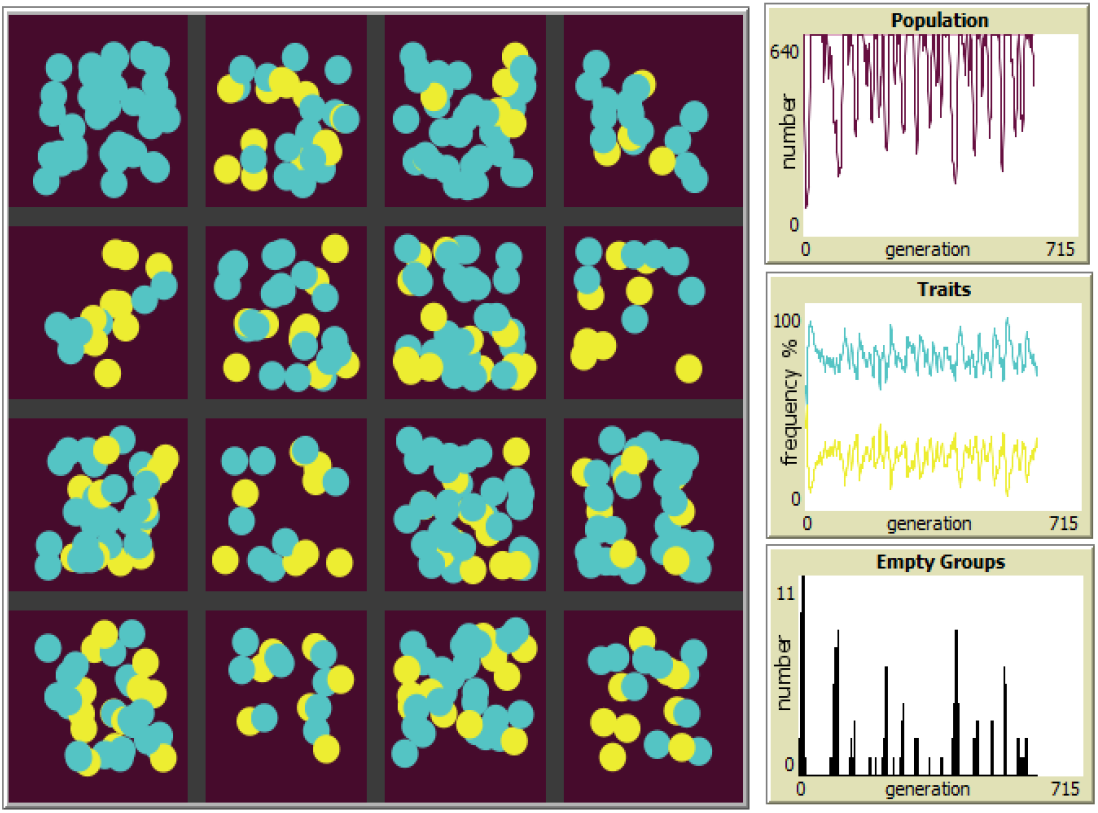
“Turbulent” (chaotic?) steady state over 600 generations as simulation outcome for larger groups and with greater differences between selection pressures for **C** and **D**. Cooperators **C** blue, defectors **D** yellow. 16 groups with a maximum of 50 individuals each. Offspring per survivor N=1; Mutation rate m=0.02; Mortality for C 25%, for D 15%. “Traits” diagram shows strong fluctuations of **C** and **D**, and also the population size and number of empty groups vary greatly, almost to extinction.

Figure 9 summarizes the simulation results for various parameters. The intrinsic base mortality is 20% in all simulations. In order to take into account the additional costs for the cooperative trait, I have measured series with an additional mortality of −5 to 25% for ***C***. The intrinsic mortality rates for ***C*** thus rank from 15 to 45%, that of ***D*** is always 20%.

**Figure 9:**
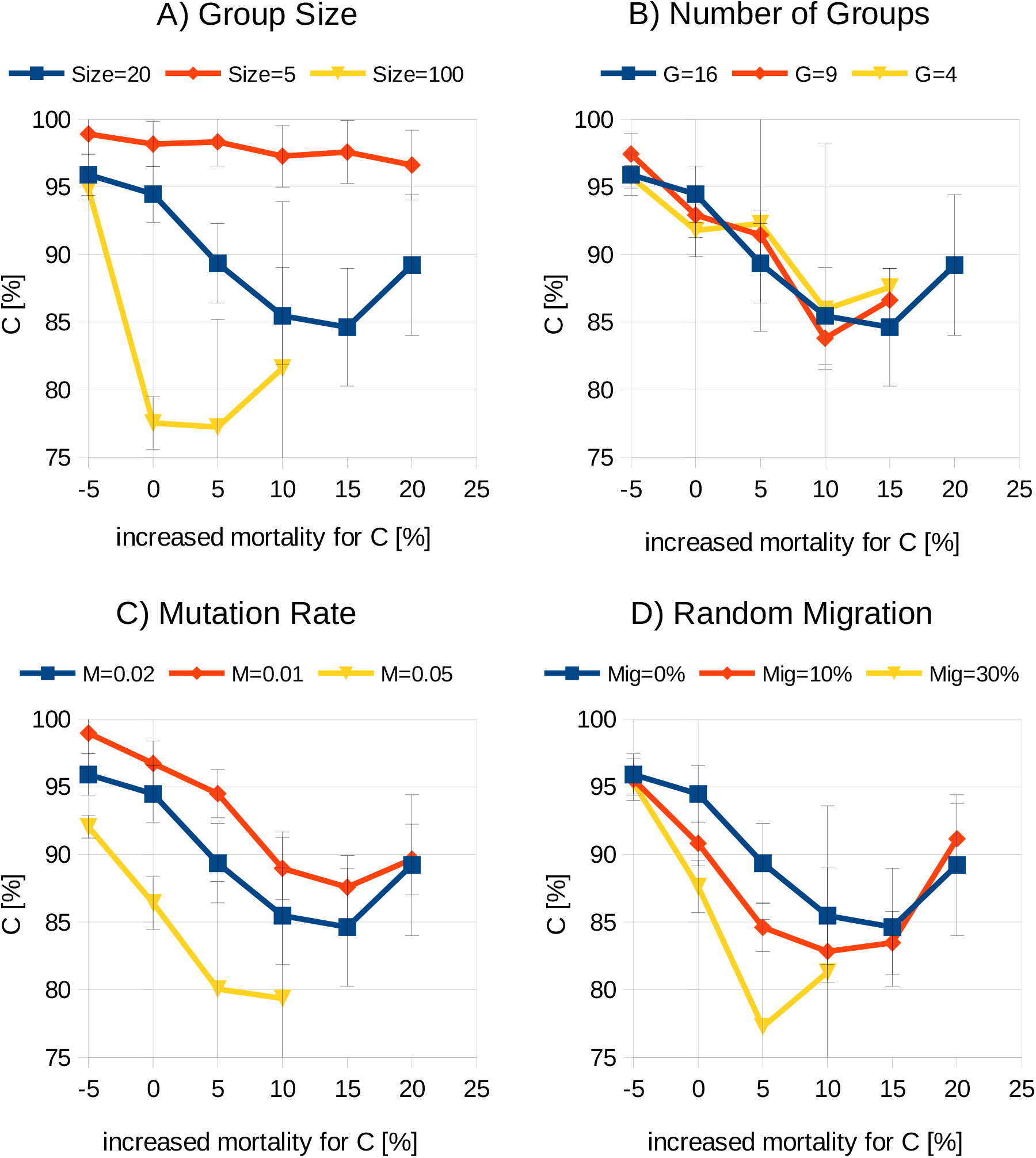
Possible mortality rates (=possible costs) for **C** and their frequencies in the population for (A) various groups sizes, (B) numbers of groups, (C) mutation and (D) random migration rates. The mortality for trait D was always 20%, the “increased mortality for C” is relative to this 20 %. The blue lines are the reference system given in Fig 7. Error bars show the standard deviation during 100 generations (after 100 initial generations to settle in). Note that if a graph ends, for example, at 10% “increased mortality for C”, it means that at 15% the whole population died out in ten out of ten attempts before the 200th generation.

The simulations show that, under suitable circumstances, the grouping effect can compensate for up to 20% higher mortality, i.e. twice as high mortality for ***C*** compared to ***D***.

In addition to the possible mortality rates, Fig. 9 shows the proportions of ***C*** in the population for various parameters. It can be seen that there are fewer ***C*** and therefore more ***D*** in the settled population if the group sizes, mutation rates and/or random migration rates are larger. Conversely, there are fewer ***D*** in the population when the groups are small, the mutation is low, and there is little migration. The number of groups, on the other hand, shows no significant influence on the ratio of ***C*** and ***D***.

In addition to this simple asexual model, I have also examined simulations with sexual reproduction, with and without generation overlap, and others with graduated traits instead of the binary C vs. D. In all of them, dynamic equilibria as described above were found over wide value ranges.

### Wilson’s trait group model and the grouping effect

Finally, I want to come back to Wilson’s trait group model and its review in Maynard Smith & Szathmáry (1996). In Tab. 1, I demonstrate how this stochastic grouping effect affects and solves the example of the trait group model shown in Fig. 1. With the same first gene pool and the same group division, as shown in Fig. 1, this effect results in three cooperators ***C*** (altruists) more and three defectors ***D*** (egoists) less in the second gene pool (13C12D) than in the first (10C15D). These are two ***C*** fewer and two ***D*** more than shown in the fictitious second gene pool of Fig. 1 with 15C10D, but nevertheless cooperation has increased significantly! This just happens at random, without synergistic effects, and also without competition between groups.

**Table 1:** Stochastic redistribution of altruistic (C) and egoistic (D) replicators through group formation using the example of the scheme shown in Fig. 1. Again, the assumption is that every altruist (C) makes the contribution b=1 to the group benefit, egoists contribute nothing (b=0) and the sum of a group’s benefits directly determines its survival rate. As result, the gene pool changes from 10C15D to 13C12D, so there are three more altruists in the second gene pool than in the first, and accordingly three less egoists.

| First gene pool | Groups | Second gene pool |
| --- | --- | --- |
| **10C15D**<br>i.e. the overall benefit is...<br>$B=10$<br>... divided in ...<br>$B_C=4$<br>$B_D=6$ | 1C4D: $B_{C1}=1/5$ and $B_{D1}=4/5$ | The benefit $B=10$ is now divided in ...<br>$B_C=\sum B_{Cn}=26/5$<br>$B_D=\sum B_{Dn}=24/5$<br>If the trait benefit correlates with survival, then the proportional replication up to the initial populations ($N = 25$) results in ...<br>**13C12D** |
| | 2C3D: $B_{C2}=4/5$ and $B_{D2}=6/5$ | |
| | 2C3D: $B_{C3}=4/5$ and $B_{D3}=6/5$ | |
| | 1C4D: $B_{C4}=1/5$ and $B_{D4}=4/5$ | |
| | 4C1D: $B_{C5}=16/5$ and $B_{D5}=4/5$ | |

## Conclusion

In this study I show that dividing a population of cooperators and defectors into groups strengthens the cooperators through purely stochastic effects which appear in the distribution of the beneficial contributions made by the cooperators. The key factor behind this grouping effect is the composition of the groups: the greater the dispersion of ***C***:***D*** ratios among the groups, the greater the effect. This advantages can to a considerable extent compensate the selection pressure that may be associated with the cooperative trait.

Despite this grouping effect, each group of replicators can still be unstable and quickly disappear on its own, especially when there is a high proportion of defectors or the selection pressure against cooperators is very high. However, if low-individual groups are provided with emigrating offspring from prospering groups, which mainly consist of cooperators, a dynamic equilibrium will usually be established between the groups, and thus the population regulates itself. In other words, no further mechanism like reciprocity or kin selection is required in this concept to explain the emergence and maintenance of a stable cooperative trait in the population.

Alarm signals within animal groups, for example, could be easily explained with this grouping effect, since a signal like “Watch out, predator!” has a direct influence on the group’s mortality. As long as the signaling does not exceed a certain risk for the emitter, it will be established and maintained in the population. In this context, groups can also go beyond species boundaries, so that the grouping effect can also provide an explanation for mutualism.

Finally, how is this grouping effect to be classified in terms of evolutionary theory? Groups are essential for the described effect. Grouping causes a stochastic redistribution of the benefit payoffs to the advantage of the cooperative trait and consequently favors the survival and reproduction chances of cooperators. This change of odds is an emergent phenomenon of any division of a population of cooperators and defectors in different groups. Neither competition between the groups nor non-random sorting or synergistic fitness effects are required to increase the frequency of cooperative replicators. The grouping effect is sufficient that cooperators contribute disproportionately to the next generation and ultimately the cooperative traits have an advantage over the selfish ones in the population.

## Acknowledgments

This work was carried out privately, independently and without institutional affiliation or funding. My heartfelt thanks go to Karin and our daughters Salima and Lara for their love, support, discussions and for their patience when I repeatedly disappeared into my world of thought experiments and simulated realities.

Supplemental material like the online versions of my simulation programs can be found at https://www.vinckensteiner.com/ksteiner

## Notes

http://www.vinckensteiner.com/ksteiner/

### Competing Interest Statement

The authors have declared no competing interest.

## References

Maynard Smith, J. & Szathmáry, E. The Major Transitions in Evolution (Freeman, Oxford, 1995).

Nowak, M. A. (2012). Evolving cooperation. J. Theor. Biol. 299: 1-8. PMID: 22281519. DOI: 10.1016/j.jtbi.2012.01.014

Traulsen, A., & Nowak, M. A. (2006). Evolution of cooperation by multilevel selection. Proceedings of the National Academy of Sciences, 103(29), 10952–10955.

Wilensky, U. (1999). NetLogo. http://ccl.northwestern.edu/netlogo/. Center for Connected Learning and Computer-Based Modeling, Northwestern University, Evanston, IL.

Wilson, D. S. (1975). A theory of group selection. Proceedings of the national academy of sciences, 72(1), 143–146.

Wilson, D. S., & Wilson, E. O. (2008). Evolution “for the Good of the Group”: The process known as group selection was once accepted unthinkingly, then was widely discredited; it’s time for a more discriminating assessment. American Scientist, 96(5), 380–389.

